# Conidial melanin of the human pathogenic fungus *Aspergillus fumigatus* disrupts cell autonomous defenses in amoebae

**DOI:** 10.1101/730879

**Authors:** Iuliia Ferling, Joe Dan Dunn, Alexander Ferling, Thierry Soldati, Falk Hillmann

## Abstract

The human pathogenic fungus *Aspergillus fumigatus* is a ubiquitous saprophyte that causes fatal infections in immunocompromised individuals. Following inhalation, conidia are ingested by innate immune cells and can arrest phagolysosome maturation. How such general virulence traits could have been selected for in natural environments is unknown. Here, we used the model amoeba *Dictyostelium discoideum* to follow the antagonistic interaction of *A. fumigatus* conidia with environmental phagocytes in real time. We found that conidia covered with the green pigment 1,8-dihydroxynaphthalene-(DHN)-melanin were internalized at far lower rates when compared to those lacking the pigment, despite high rates of initial attachment. Immediately after uptake of the fungal conidia, nascent phagosomes were formed through sequential membrane fusion and fission events. Using single-cell assays supported by a computational model integrating the differential dynamics of internalization and phagolysosome maturation, we could show that acidification of phagolysosomes was transient and was followed by neutralization and, finally, exocytosis of the conidium. For unpigmented conidia, the cycle was completed in less than 1 h, while the process was delayed for conidia covered with DHN-melanin. At later stages of infection, damage to infected phagocytes triggered the ESCRT membrane repair machinery, whose recruitment was also attenuated by DHN-melanin, favoring prolonged persistence and the establishment of an intracellular germination niche in this environmental phagocyte. Increased exposure of DHN-melanin on the conidial surface also improved fungal survival when confronted with the fungivorous predator *Protostelium aurantium*, demonstrating its universal antiphagocytic properties.

## Introduction

The ubiquitous filamentous fungus *Aspergillus fumigatus* is most commonly found in the soil or on decaying organic matter and randomly infects immunocompromised individuals after inhalation of its conidia (Brakhage and Langfelder, 2002). Over 200,000 life-threatening infections are caused by *A. fumigatus* annually, with mortality rates of infected individuals ranging from 30-90% (Brown et al., 2012, Bongomin et al., 2017). Poor diagnosis, often rapid disease progression and gaps in our understanding of the early stages of infection are currently limiting therapeutic options.

The green-greyish conidial pigment 1,8-dihydroxynaphthalene (DHN)-melanin is among the first microbe-associated molecular patterns that the host encounters during infection. It has been shown that the presence of DHN-melanin interferes with conidial uptake and processing in mammalian phagocytes and can inhibit apoptosis in endothelial lung cells (Thywissen et al., 2011, Slesiona et al., 2012, Heinekamp et al., 2012, Volling et al., 2011, Amin et al., 2014, Akoumianaki et al., 2016, Jahn et al., 2002). Myeloid and endothelial cells of the lung recognize DHN-melanin itself via the C-type lectin receptor MelLec, which plays an important role in the protective antifungal immunity of both mice and humans (Stappers et al., 2018). The processing of conidia by phagocytic cells is crucial to understand, as these cell types are involved in the defense and also the dissemination of the fungus. Recent experiments with macrophages showed that melanized conidia of *A. fumigatus* interfere with phagosome acidification by preventing the formation of lipid rafts that are essential for v-ATPase proton pump assembly (Schmidt et al., 2019). Upon germination, the DHN-melanin layer is lost, exposing chitin, glycoproteins, and β-1,3-glucan, whose exposure facilitates recognition, phagocytic uptake and killing by immune cells (Chai et al., 2010, Luther et al., 2007). The biochemical fate of fungal melanin following swelling and germination is thus far unknown.

In contrast to commensal pathogens such as *Candida albicans*, *A. fumigatus* is considered an environmentally acquired pathogen, as it is frequently isolated from natural reservoirs and occupies a well-established niche as a decomposer of organic matter. In its natural environment the fungus is confronted with many abiotic and biotic adverse conditions such as amoebae with some of them being able to ingest and even kill *A. fumigatus* (Hillmann et al., 2015, Radosa et al., 2019a). During evolution it can thus be expected that microorganisms such as *A. fumigatus* have acquired counter defense strategies that also might explain the virulence of environmental pathogens for humans. This hypothesis was recently coined as the “Amoeboid predator-animal virulence hypothesis”. According to this hypothesis microorganisms trained their virulence through competition with microbial predators (Casadevall et al., 2019). In agreement with this hypothesis, several recent studies demonstrated that *A. fumigatus* interactions with soil amoeba such as *Acanthamoeba castellanii* or *Dictyostelium discoideum* exhibited similar outcomes to its interactions with human phagocytes (Van Waeyenberghe et al., 2013, Hillmann et al., 2015, Mattern et al., 2015).

*D. discoideum* is a professional soil phagocyte that constantly engulfs microbes for food and thus has to protect itself from potentially harmful intracellular pathogens (Cosson and Soldati, 2008, Dunn et al., 2018). To avoid infection, the phagocytic host has to eliminate pathogens by forming a functional phagosome before they can escape or establish a survival niche. After engulfment, the pathogen is trapped inside the nascent phagosome, which is mainly derived from the plasma membrane. Initially, it lacks any microbicidal capacity that is essential for pathogen control. By a sequence of membrane fusion and fission events, the phagosome acquires its full range of antimicrobial and degradative features. This conversion is known as phagosome maturation, during which the compartment undergoes consecutive fusion with early and late endosomes and lysosomes (Flannagan et al., 2009). The final step of phagosome maturation is its resolution, during which the phagosomal content becomes recycled, and indigestible material is exocytosed (see (Dunn et al., 2018) for review).

The majority of intracellular pathogens reside in a vesicular compartment, where they hijack the defense machinery of the cell to get access to host resources, but some bacteria have evolved an arsenal of strategies to invade the host cytosol by utilizing pore-forming toxins, phospholipases or effector-delivery systems, Examples include *Listeria* and *Shigella* which launch an early escape, while *Mycobacteria* and *Salmonella* execute a partial or delayed escape from the phagosome (Friedrich et al., 2012). In contrast, the human pathogenic fungus *Cryptococcus neoformans* escapes from *D. discoideum to the extracellular space* by both WASH-mediated constitutive exocytosis and vomocytosis (Watkins et al., 2018).

Professional phagocytes developed various methods to combat intracellular microorganisms establishing a toxic, bactericidal environment inside the phagosome and using a combination of cytosolic machineries, such as ESCRT and autophagy, to restrict the pathogens. Plenty of relevant environmental pathogens have co-evolved with their hosts to overcome the defenses of the host cell, which allows them to proliferate or achieve a state of latency. As conidia of *A. fumigatus* were shown to interfere with phagolysosome maturation in phagocytes from mammalian hosts, we have used two model amoeba to assess whether this strategy of the fungus may have even broader host specificity and thus, could provide a selective advantage for the fungus in its natural environment.

## Results

### 1,8-DHN-melanin delays phagocytic uptake and phagolysosome maturation

To initiate phagocytosis, host receptors engage with ligands exposed on the surface of *A. fumigatus* conidia. This association with its ligand initiates signaling pathways that cause the extension of a lamellipodium, which surrounds the particle and generates the nascent phagosome. The surface layer of wildtype fungal conidia consists of α-1,3-glucan covered by DHN-melanin and a hydrophobic, proteinaceous rodlet layer. The surface of conidia of the melanin-deficient mutant Δ*pksP* is composed of the rodlet layer, glycoproteins, and chitin (Valsecchi et al., 2018).

We first analyzed infection outcomes after co-incubation of *D. discoideum* with wild-type and *pksP* mutant conidia of *A. fumigatus*. After 1 h of co-incubation we found that *D. discoideum* amoebae had ingested 63% of the melanin-deficient Δ*pksP* conidia, but only 20% of the wild-type conidia (Figure 1A-B). The phagocytic efficiencies determined for wild-type and Δ*pksP* conidia were lower and higher than the ones for inert silica particles, respectively (Figure 1C and S1A). Melanin ghosts obtained after harsh chemical treatment of wild-type conidia were rarely taken up by the amoeba. However, these empty shells of melanin would readily associate with the amoeba cell wall, covering the entire surface (Figure 1A+B and Figure S1A). Collectively, our results suggested that DHN-melanin might have an impact on the uptake process of conidia. This conclusion was further supported by an experiment with the DHN-melanin monomer 1,8-dihydroxynaphthalene, which also repressed phagocytosis of beads in a dose-dependent manner (Figure 1D).

**Fig 1.**
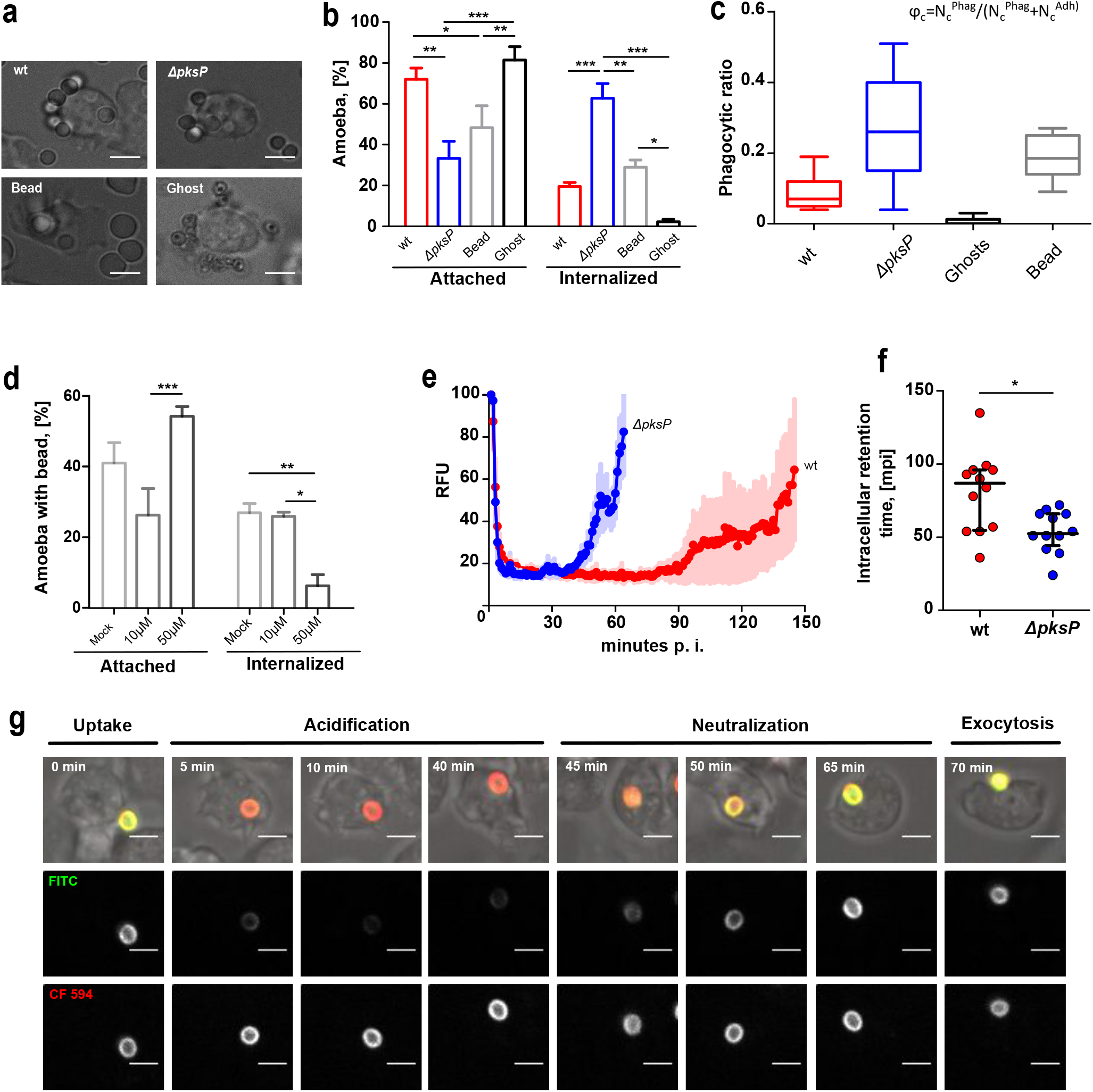
Phagocytosis of *Aspergillus fumigatus* conidia by *Dictyostelium discoideum*. **a** Resting conidia of the wild type (wt) or the melanin-deficient strain of *A. fumigatus* (Δ*pksP*) were added to *D. discoideum* at MOIs of 5. Silica beads (Bead) and melanin ghosts (Ghost) were added to the amoebae at the same MOI. Images were captured after 1 h of co-incubation. The scale bar is 5 μm. **b** Cells with phagocytic and attachment events were quantified from images captured at 1 h p. i. The bars represent the mean and SEM from three independent experiments, with n=100 for each experiment. Statistical differences were calculated with a Bonferroni posttest after a two-way ANOVA with asterisks indicating significance (*p<0.05; **p<0.01; ***p<0.001). **c** Phagocytic ratio for *A. fumigatus* conidia, silica beads and melanin ghosts. **d** Wild type amoeba were exposed to silicon beads in the presence of 10 or 50 μM of 1,8-DHN. Imaging and quantification were carried out as in b. **e-g** Amoebae were infected with resting conidia of the wild type or the Δ*pksP* strain pre-stained with the pH-sensitive fluorophore (FITC) and the reference fluorophore (CF594) for realtime measurements of acidification and residence time in the amoeba. **e** Timeline of FITC derived fluorescence intensity indicating pH variations at the conidial surface during phagocytosis. **f** The intracellular retention time of conidia inside of *D. discoideum*. Statistical differences were calculated with a t-test. **g** Time-lapse illustration of major steps during the phagocytic cycle for resting conidia of the Δ*pksP* mutant.

We were further interested in the intracellular fate of the conidia and frequently observed conidial exocytosis. We thus followed the infection process at the single-cell level and monitored the time of intracellular transit of conidia in *D. discoideum*. Conidia were stained with FITC (green) and with CF594 (red) for the normalization of signal intensity (Figure S1B and S2). Ratiometric calculations of the differences between the two dyes, with FITC responding to changes in pH, allowed us to track the phagosomal pH dynamics for conidia over the entire intracellular period (Figure 1E). These measurements demonstrated that both wildtype and Δ*pksP* mutant conidia underwent rapid acidification followed by neutralization and subsequent exocytosis. This phagosomal processing was previously reported to also occur with inert particles, which complete the process of acidification, neutralization and exocytosis within 1 h (Gopaldass et al., 2012). Phagosome maturation and exocytosis of resting melanin-deficient conidia were completed at a time scale resembling that of of inert particles (Figure 1E). Notably, wild-type conidia resided significantly longer inside phagolysosome than Δ*pksP* conidia (Figure 1F, Figure S1B), which suggested interference at single or multiple stages of the phagosome maturation process.

### Functional v-ATPase is trafficked to *A. fumigatus* containing phagosomes

Proton transport into intracellular organelles is mainly accomplished by the vacuolar-ATPase (v-ATPase). This enzyme complex is composed of two multi-subunit domains, which together pump H^+^ into the lumen in an ATP-dependent manner. In *D. discoideum* the v-ATPase is mainly localized at membranes of the contractile vacuole, an osmoregulatory organelle, and at the membranes of endosomes to generate their acidic lumenal environment (Liu and Clarke, 1996). To visualize the real-time distribution of the enzyme complex, we employed *D. discoideum* strains expressing fusions of the v-ATPase membrane channel subunit VatM and the cytosolic domain VatB with green and red fluorescent proteins, respectively (Clarke et al., 2002). A combination of these two marker proteins previously revealed the principal route of delivery of the v-ATPase to phagosomes (Clarke et al., 2010).

Live, single-cell imaging of FITC-stained conidia after uptake by VatB-RFP-expressing amoeba demonstrated the correlation between v-ATPase recruitment and acidification. Fast acidification and v-ATPase trafficking to the surface of the phagosome was followed by its retrieval and subsequent neutralization of the phagosomal lumen, with conidial exocytosis as the final step (Figure 2A). As expected from the previous experiments, the acidification kinetics for wild-type and Δ*pksP* conidia varied significantly. Both fungal strains triggered acidification within minutes, but amoebae infected with wild-type-conidia were delayed in reaching the minimum and maximum pHs (Figure 2B). Also, following VatB-RFP retrieval, phagosomes took significantly longer to reach pH 6 again when infected with DHN-melanin-covered wild-type conidia when compared to Δ*pksP* conidia-containing phagosomes (Figure 2C).

**Fig 2.**
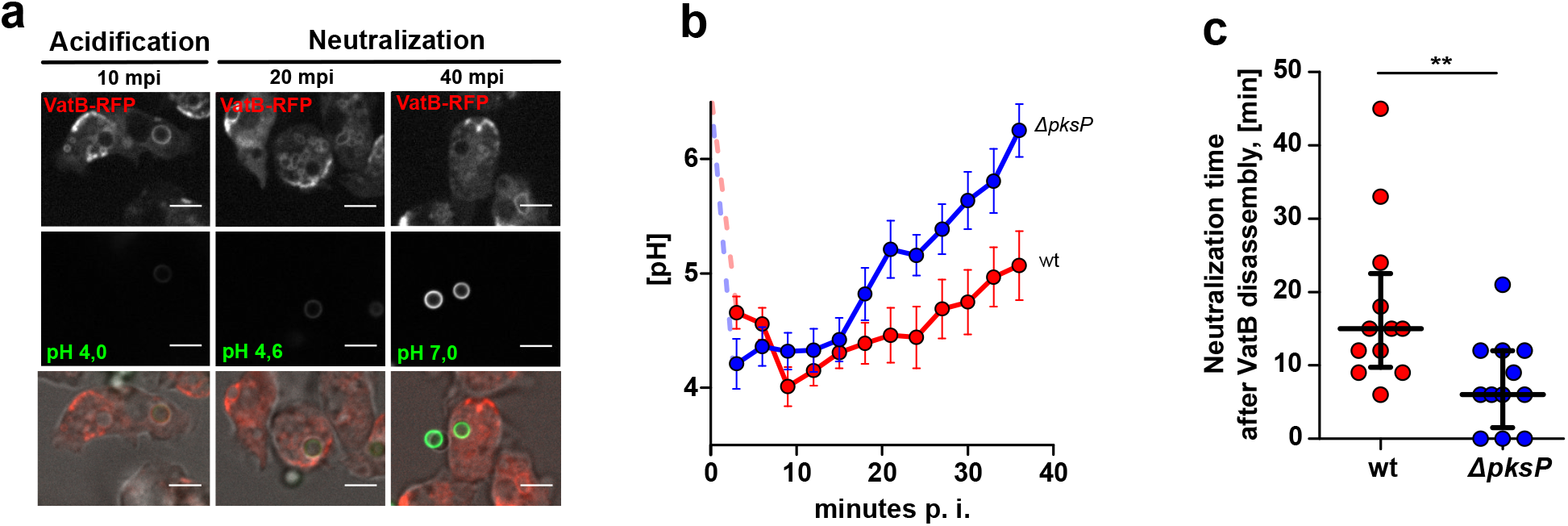
v-ATPase assembly and phagosomal acidification during conidial infection. **a** Representative images of different stages of phagosome maturation in VatB-RFP expressing cells infected with FITC stained Δ*pksP* conidia. Scale bar is 5μm. **b** Kinetics of phagosomal pH in VatB-RFP expressing cells infected with resting conidia. Twelve independent movies were taken for each fungal strain. Dots and error bars indicate the Mean and SE respectively. **c** Phagosomal neutralization time (pH 6.0) after VatB disassembly from the phagosomal surface. Statistical differences were calculated with a Student’s t-test with P=0.0066

### A simulation-based prediction of maturation dynamics in long-term confrontations

The different dynamics of uptake, acidification and exocytosis for both strains determined from single-cell observations were used in a Monte-Carlo simulation to predict the outcome of long-term confrontations (Figure 3A and B). The simulation was executed with the following experimentally determined parameters for wild type and Δ*pksP* conidia, respectively: conidial uptake probability of 20 and 63%, acidic time spans of 52 and 32 min, and exocytosis at 87 and 53 min after uptake. Assuming an amoeba population of 10^4^ cells infected at an MOI of 10 (10^5^ conidia in total), the computational simulation predicted that the different rates of uptake and phagosome maturation for wildtype and melanin-deficient conidia would yield significantly different acidification patterns across the infected amoeba populations. Infections with melanized *vs*. non-melanized conidia yielded 100% *vs*. 82% of acidified phagolysosomes after 30 mpi, respectively, while 36% of phagolysosomes containing wild-type conidia and 48% of those containing Δ*pksP* conidia-were acidified after five hours of co-incubation. As the period between uptake and acidification of the phagosome was too short to allow for accurate measurement, we set this value to zero, which may have influenced the precision of the computational model. Nevertheless, the time frame for the trafficking of the v-ATPase (Figure 3C-F), as well as the fact that nearly all v-ATPase-containing phagosomes were also acidic (Figure S3) confirmed the Monte-Carlo model prediction with reasonable accuracy and further supports the finding that phagosome maturation is delayed by the presence of DHN-melanin.

**Fig 3.**
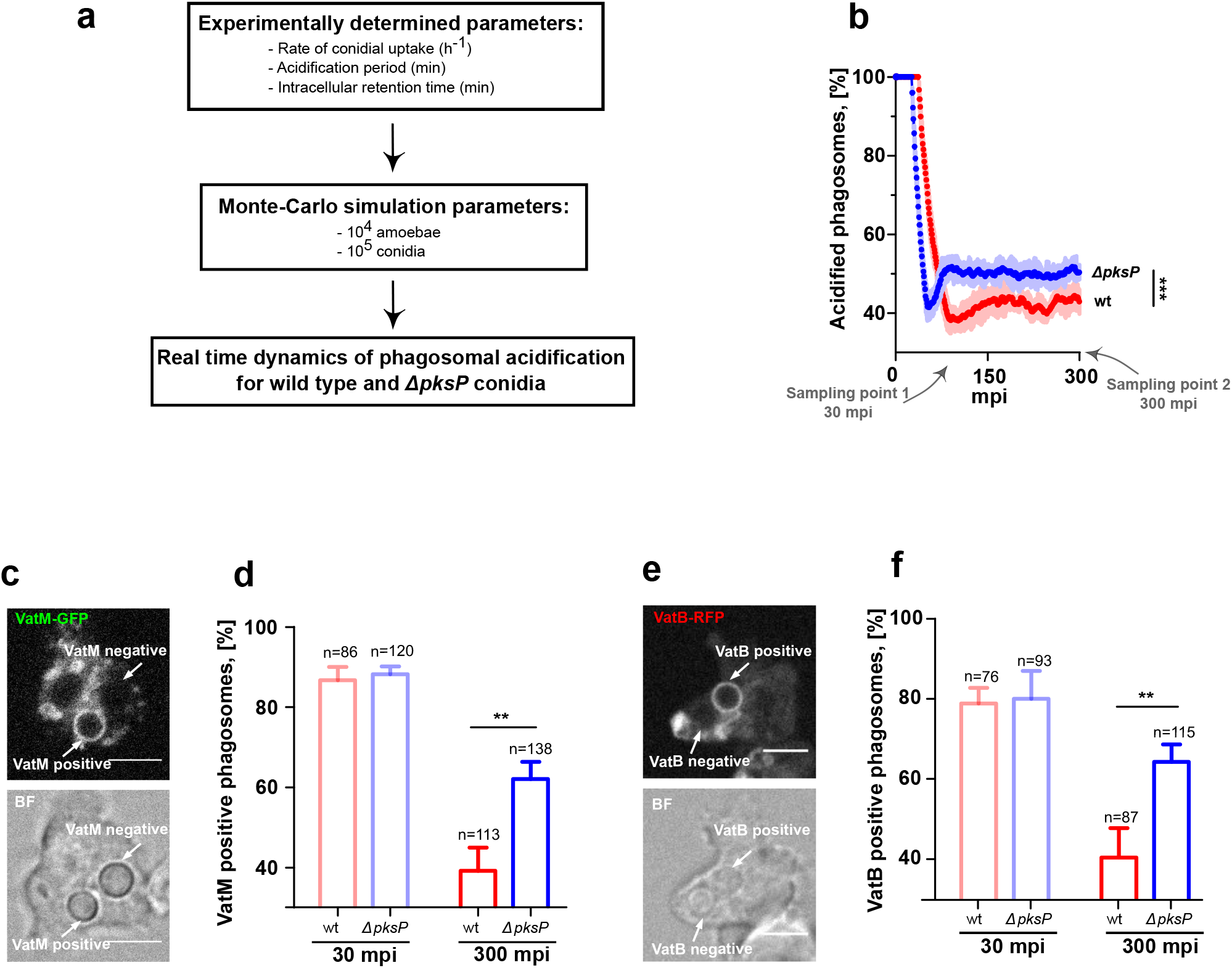
Quantification of number acidified phagosomes in the whole amoeba population. **a** Setup for the computational simulation of population dynamics based on experimentally determined parameters in single-cell analyses **b** Percentage of acidic phagosomes in the population of 10^4^ amoeba infected with wild type and Δ*pksp* conidia at an MOI of 10. Statistical differences were calculated with a Bonferroni post-hoc test after a two-way ANOVA (p<0.0001). **c, e** Representative images of VatM-GFP localization after 30 min p. i. (**c**) and VatB-RFP localization after 300 min p.i. (**d**) The scale bar is 5 μm. **d, e** Percentage of VatM-GFP (**e**) and VatB-RFP (**f**) positive phagosomes after 30 and 300 min p. i. Experiments were performed in 3 biological replicates. Statistical differences were calculated with t-test with P=0.0053 and P=0.0085 for (**d**) and (**f**), respectively.

### ROS generation coincides with damage to *A*. fumigatus-containing phagolysosomes

NADPH oxidase (NOX) is heteromultimeric, membrane bound complex that produces intraphagosomal ROS. The enzyme also plays an important role during *A. fumigatus* infection in humans (reviewed in (Hogan and Wheeler, 2014)) *D. discoideum* encodes three NOX catalytic subunits, i.e., *noxA-C*, with NoxA and B being homologues of the mammalian gp91^phox^ subunit. A single gene, *cybA*, encodes the only *D. discoideum* homologue of the p22^phox^ subunit of the mammalian NADPH oxidase (Lardy et al., 2005, Dunn et al., 2018). With wild type conidia we detected CybA at the phagosome only after 1 h of infection (Figure 4). Further, highly acidic phagosomes were generally devoid of CybA (Figure 4A). At this time point, the phagosomal pH was still highly variable among different phagosomes even within single cells, presumably due to asynchronous internalization (Fig. 4B). After 2 h of infection, 81 % of all phagosomes were positive for ROS, but even in amoebae lacking all three *nox* genes, ROS production was detected in approximately 50 % of all phagosomes, demonstrating that the NOX proteins are a substantial but not only source of ROS in phagosomes (Figure 4C and Figure S4). At this stage of infection, CybA-positive phagosomes containing the melanin-deficient Δ*pksP* conidia displayed an average pH of around 6.2, while the pH of CybA-positive phagosomes containing wild-type conidia was approximately 7.2 (Figure 4D). This relatively high pH may be attributed either to higher NADPH oxidase activity neutralizing the phagosome more effectively through increased formation of ·O_2_^−^ or, more likely, to a leakage of protons from the phagosome mediated by damage to the phagosomal membrane.

**Fig 4.**
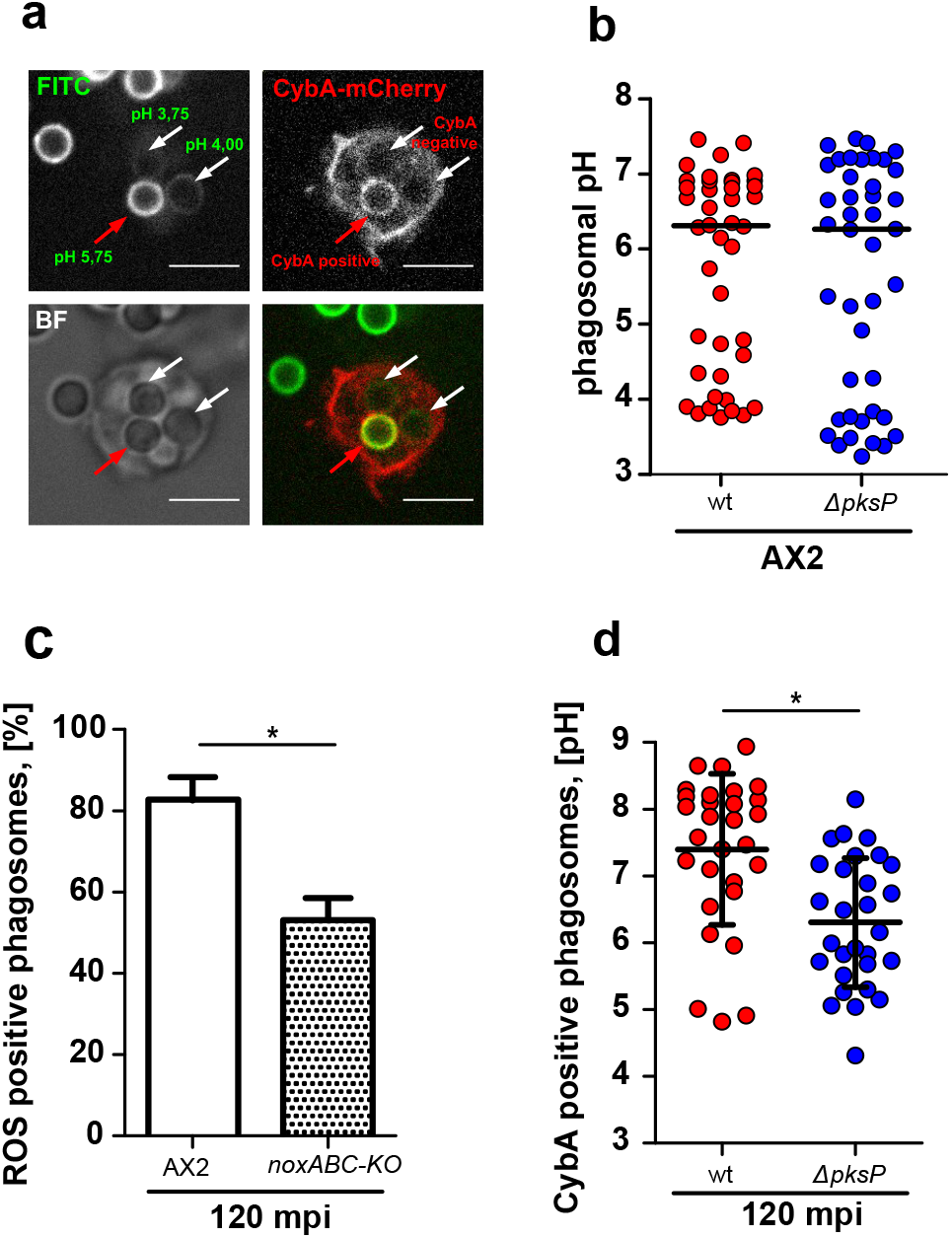
NADPH oxidase trafficking follows phagosome neutralization. **a** Representative image of *D. discoideum* expressing CybA-mCherry after 1 h of infection with FITC stained conidia of the wild type to assess the phagosomal pH and NADPH oxidase activity simultaneously. The scale bar is 5 μm. **b** FITC based pH measurement of phagosomes 1 h p. i. Experiments were performed in 3 biological replicates and statistical differences were calculated with a Bonferroni post-hoc test after a two-way ANOVA. **c** DHE based quantification of ROS in phagosomes of wild type *D. discoideum* and mutant lacking all three nox genes. Experiments were performed in 3 biological replicates. Statistical differences were calculated with the t-test. **d** FITC based pH measurement of infected CybA-mCherry-positive phagosomes 2 h p. i. Experiments were performed in 3 biological replicates. Statistical differences were calculated with a t-test.

### Lysosome fusion indicates damage to *A*. fumigatus-containing phagosomes

Proper phagosome maturation involves the fusion of early/late phagosomes with lysosomes, which load proteolytic enzymes for digestion of the phagolysosomal content. To monitor lysosomes and their fusion with phagosomes, the lysosomes of amoebae were loaded with fluorescently labelled 70-kDa-dextran prior to infection with conidia. When loaded lysosomes fused to conidia-containing phagosome dextrans were visible as a ring around the conidia (Figure 5A). By measuring the normalized integrated density of these rings, we concluded that the phagolysosome fusion was equally effective for melanized and non-melanized conidia (Figure 5B). We substantiated this data by analyzing vacuolin, a postlysosomal marker that represents a functional homologue of the metazoan lipid-raft microdomain chaperon flotillin (Bosmani et al., 2019). In agreement with the data obtained for dextran accumulation, vacuolin gradually accumulated in the membrane of both phagolysosomes containing either wildtype- or Δ*pksP*-conidia (Figure S5). Collectively, these data suggested that the lysosomal fusion is not inhibited.

**Fig 5.**
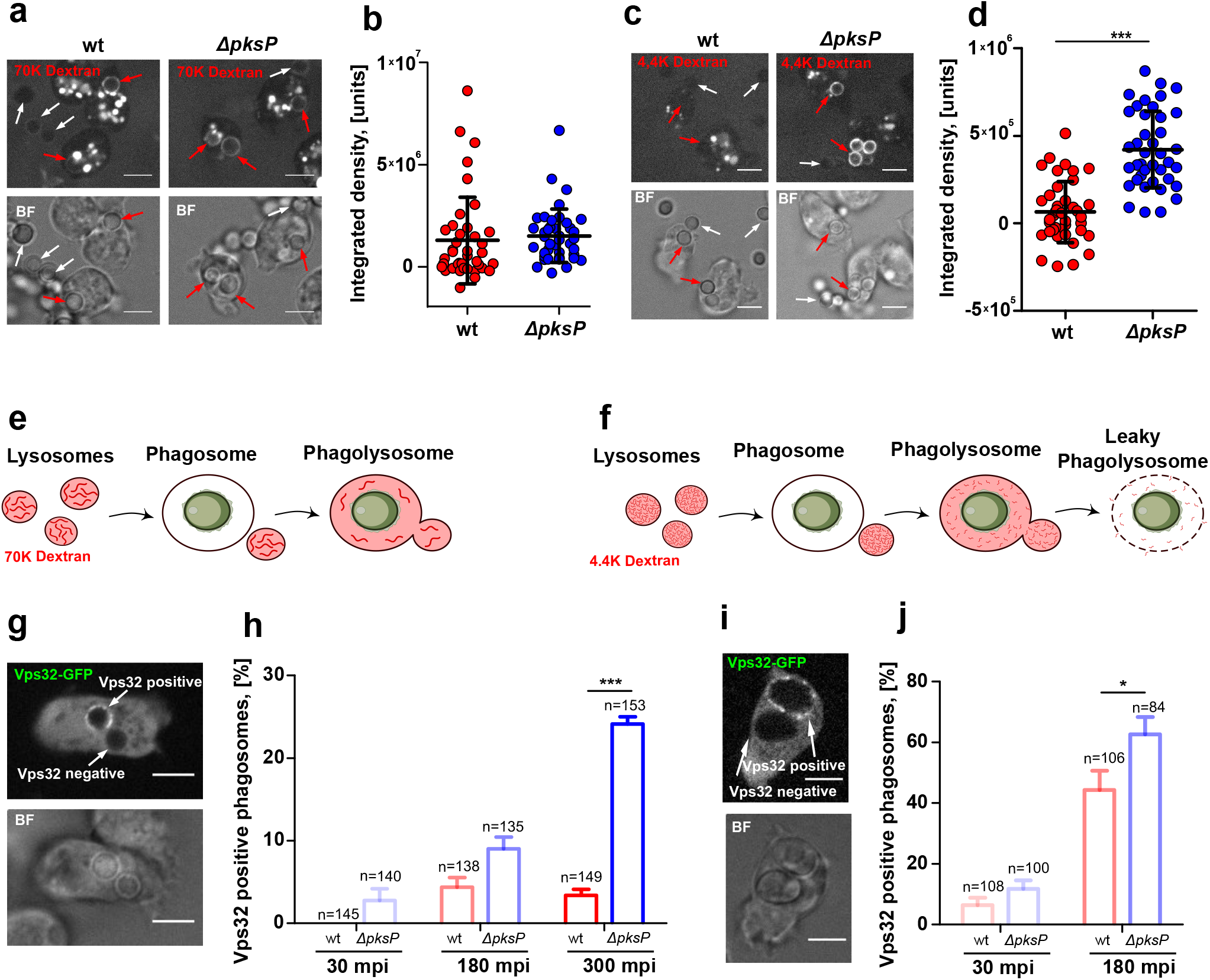
Dextran leakage and Vps32 recruitment at conidia containing phagolysosomes. **a, c** Cells of *D. discoideum* were loaded with RITC-dextran at a molecular weight of 70,000 Da (**a**) or 4.400 Da (**c**) and subsequently infected with *A. fumigatus* conidia. Images were captured after 300 mpi. Internalized conidia and free conidia are indicated by red and white arrows, respectively. **b, d** Quantification of RITC fluorescence of the two dextrans (B, 70,000 and C, 4,400) as integrated density in conidia containing phagosomes. Values were normalized by background substraction of free conidia. Images were captured after 300 mpi. Data are based on 3 biological replicates with statistical differences calculated in a one way ANOVA with p<0.0001. **e, f** Schematic representation of size discriminated leakage of dextran from phagolysosomes. **g, e** Vps32-GFP-expressing cells were infected with dormant (**g**) or swollen conidia (**i**) at an MOI of 10 and representative images from 180 m. p. i. are shown. **h, j** Quantification of Vps32-GFP localization to conidia containing phagosomes. Statistical differences were calculated with a Bonferroni post-hoc test after a two-way ANOVA with asterisks stating significance with *p<0.05, **p<0.01, and ***p<0.001). Scale bars are 5 μm.

Because the maturation of the phagolysosomes appeared not to be affected, we reasoned that the pH difference between CybA-positive phagosomes containing wildtype conidia and Δ*pksP* condidia resulted from differences in the integrity of the phagolysosomal membrane. Therefore, we preloaded the lysosomes of the amoeba with a dextran of the low molecular mass of 4.4 kDa. As shown in Figure 5C and 5D, amoebae infected with wild-type conidia displayed almost no rings, in contrast to the phagolysosomes of the melanin-deficient mutant, which retained the dextran (Figure 5C+D). These results suggested that wild-type conidia resided in leaky phagolysosomes contrary to the conidia of the Δ*pksP* strain (Figure 5E and F).

### The phagolysosomal ESCRT repair machinery is by DHN-melanin

Recently, it was demonstrated that disruptions of endolysosomes can be repaired by the <u>e</u>ndosomal <u>s</u>orting <u>c</u>omplex <u>r</u>equired for <u>t</u>ransport (ESCRT) machinery (Jimenez et al., 2014, Skowyra et al., 2018). In *D. discoideum*, Vps32 is a homologue of the CHP4A, B, C proteins of the ESCRT-III complex in metazoa. The protein localizes to injuries at the plasma membrane and endomembranes (López-Jiménez et al., 2018). We hypothesized that damage due to conidial infection triggers the recruitment of this complex which can be measured by the use of a Vps32-GFP expressing cell line (Figure 5G-H). Infection of this cell line with conidia of the Δ*pksP* strain triggered the recruitment of the ESCRT-III machinery to phagolysomes in a time-dependent manner, with a maximum of 25% of Vps32-positive phagolysosomes after a long-term confrontation of 5 h. In contrast, Vps32 was recruited to less than 5 % of wild-type conidia-containing phagolysosomes over the entire period. Pre-swollen conidia, which had lost their melanin coat at the the onset of germination, recruited higher levels of the Vps32 protein to the phagosome. Here, the numbers for Vps32-positive phagosomes exceeded 40 and 60 % of for the wild type and the mutant (Figure 5I+J). We further substantiated the lack of ESCRT recruitment to damaged, wild-type conidia-containing phagosomes by combining all three reporters, *i.e.*, Vps32-expressing cells preloaded with dextrans of both molecular masses were infected with either wild-type or Δ*pksP* conidia. Infections with the wild type lead to leaky phagosomes which were positive only for the 70-kDa-Dextran but devoid of the 4.4-kDa-Dextran and Vps32. Phagosomal damage, was also detected with Δ*pksP* conidia, as seen by the selective loss of the 4.4-kDa-Dextran. However, Vps32 was effectively recruited to these phagosomes, indicating active repair. This conclusion was further supported by the observation that the smaller Dextran was at least partially retained in Vps32-positive phagosomes (Figure 6).How, DHN-melanin could directly affect Vps32 recruitment is unclear, but at least in *in vitro*, synthetic DHN-melanin and melanin ghosts were more efficiently degraded by H_2_O_2_ at neutral pH, indicating that unknown degradation products of DHN-melanin could be present inside the phagolysosome (Figure S6).

**Fig 6.**
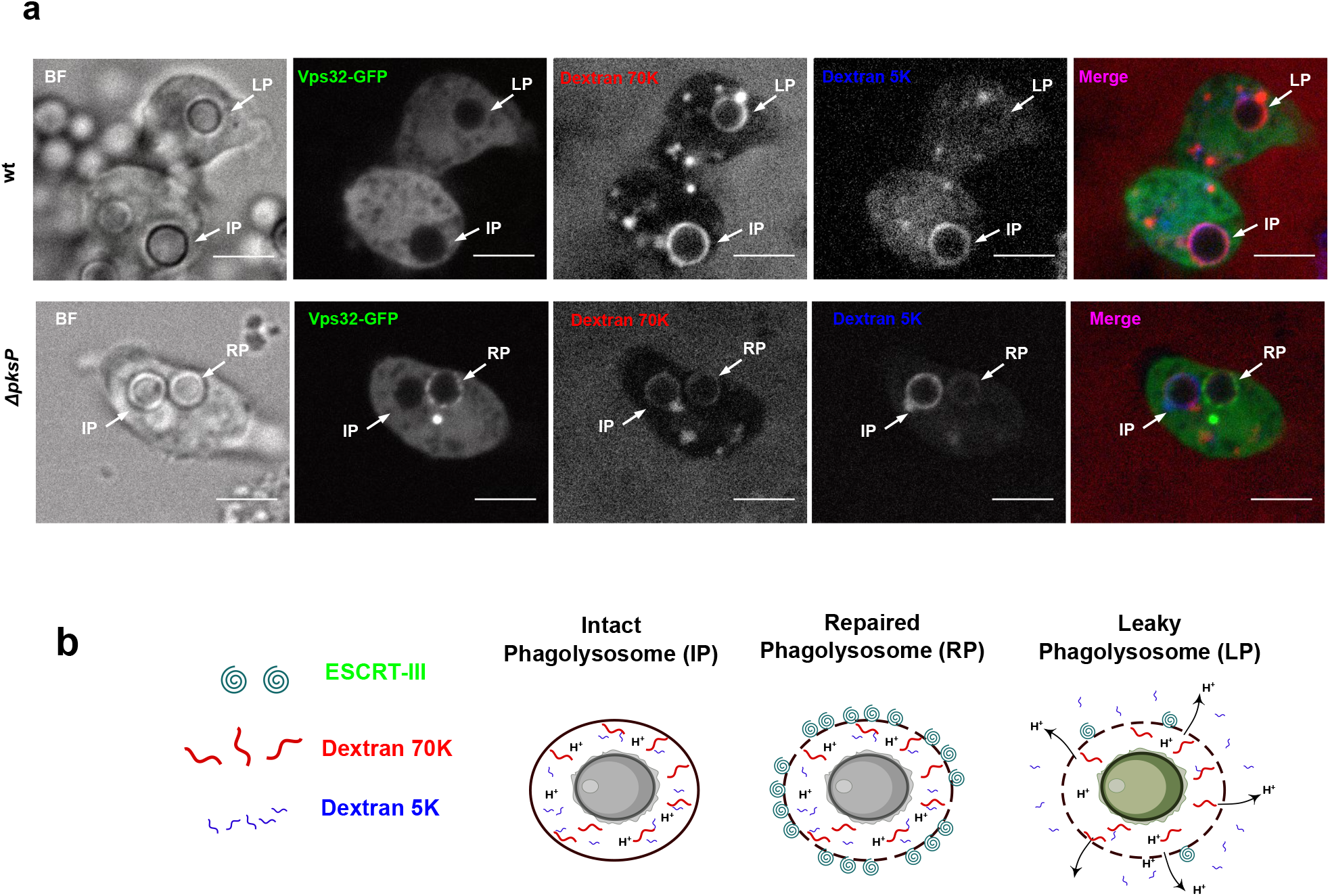
Vps32 is absent from damaged phagolysosomes containing DHN-melanized conidia. **a** Vps32-GFP-expressing cells of *D. discoideum* were first loaded with RITC-dextran of 70,000 Da and blue-dextran of 5,000 Da simultaneously and subsequently infected with dormant conidia of the wild type or Δ*pksP*. Scale bars are 5 μm. **b** Schematic illustration of the experimental results.

### DHN-melanin attenuates killing by a fungivorous amoeba

Although deformed or degraded fungal conidia after co-incubation of swollen spores with *D. discoideum* were occasionally observed after confrontation for 3 to 5 hours (Figure S7 A+B), an assay for fungal viability did not reveal any significant killing of either wild type or melanin-deficient mutant by this model phagocyte (Figure S7C). In turn, the viability of *D. discoideum* was significantly affected after 24 h of an infection by the fungus at an MOI of 0.1, but the effects were not melanin-dependent (Figure S7D). Contrary to *D. discoideum*, other amoebae are specialized mycophagous predators, with *Protostelium aurantium* being able to internalize and intracellularly digest fungal conidia (Figure S8), or invade fungal hyphae (Radosa et al., 2019b). When confronted with the fungivorous amoeba *Protostelium aurantium*, melanin-deficient conidia were killed more efficiently than wild-type conidia. Comparably higher exposure of DHN-melanin on the surface of the conidia was previously shown for fungal strains lacking the gene encoding the main conidial hydrophobin RodA (Valsecchi et al., 2018), (Figure 7A). The survival of RodA-deficient conidia was even higher than that of wild-type conidia (Figure 7B+C), suggesting that DHN-melanin can serve a protective role against this fungivorous predator.

**Fig 7.**
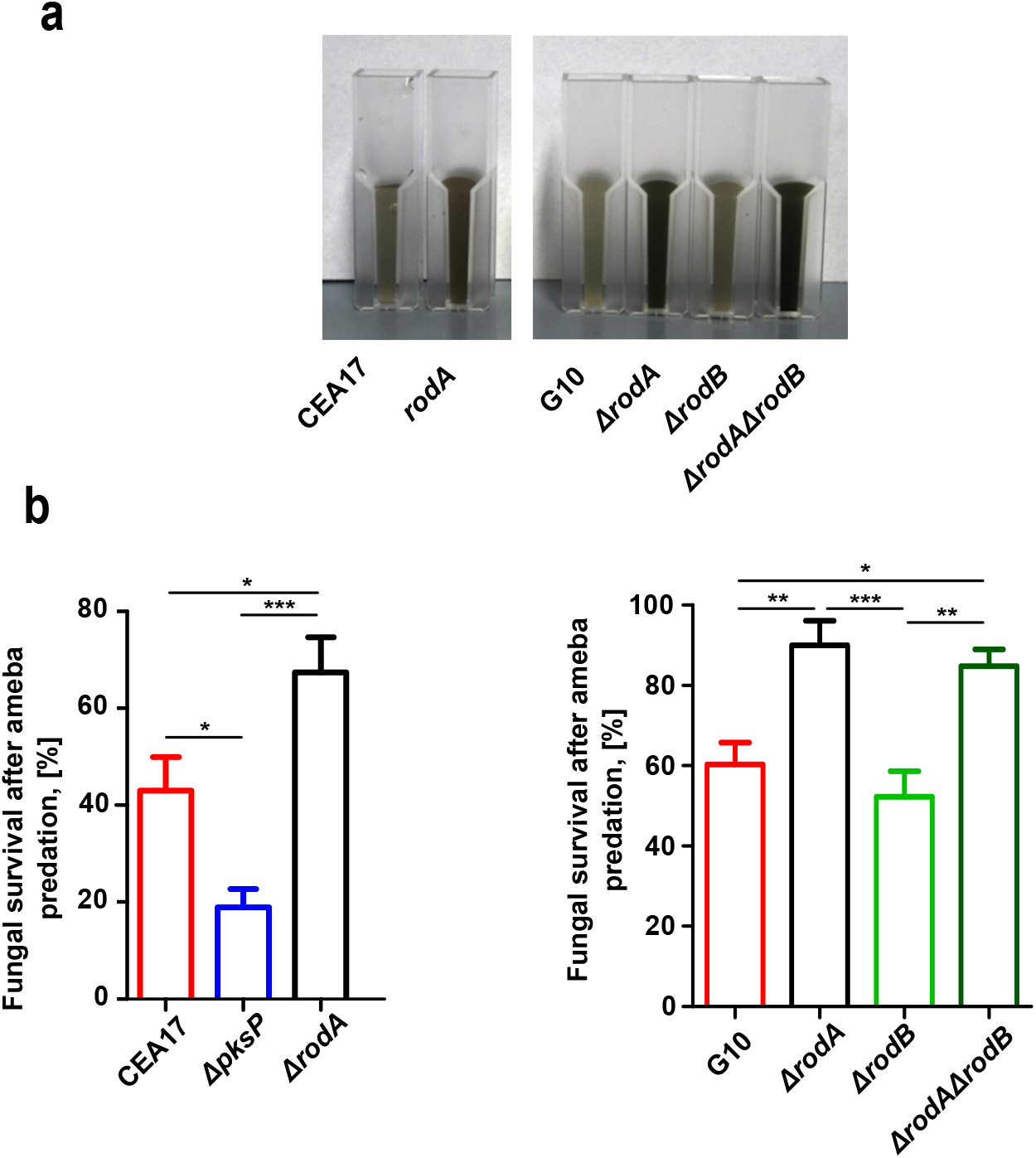
Viability of swollen conidia of *Aspergillus fumigatus* after a confrontation with the fungivorous amoeba *Protostelium aurantium. a* Suspensions of 10^9^ conidia of *A. fumigatus* strains showing different levels of DHN-melanin exposure. CEA17 and G10 represent wild type like strains, *ΔrodA, ΔrodB*, and *ΔrodAΔrodB* indicate deletion mutants for genes encoding surface hydrophobins. **b** Viability of conidia after *P.aurantium* predation. Fungal survival was determined from Resazurin based measurements of fungal growth following confrontations with the *P. aurantium* at an MOI of 10. Fungal survival is expressed as a mean and SEM from three independent experiments. Statistical differences were calculated with a Bonferroni post-test after a two-way ANOVA with significance shown as * p<0.05; ** p<0.01; *** p<0.001.

## Discussion

As the environmental reservoir of *A. fumigatus* suggests that phagocytic interference *via* DHN-melanin could also serve a protective role outside the human host, we have used two amoeba models to follow the antagonistic interaction of *A. fumigatus* conidia with amoebae in real time. Conidia covered with the green pigment DHN-melanin were internalized at far lower rates than those lacking the pigment, despite high levels of initial attachment. Similar findings previously obtained with human macrophages showed that DHN-melanin of *A. fumigatus* interferes with their phagocytosis rates (reviewed in (Heinekamp et al., 2012). Such parallels might indicate that DHN-melanin serves as a protective pigment against a wide range of phagocytic cells, which may either belong to the innate immune defense of metazoa or be distant members within the highly diverse kingdom of amoebozoa.

We provided further evidence that the first intracellular processing steps in the amoeba, v-ATPase trafficking and acidification, are only marginally affected during *A. fumigatus* infection of *D. discoideum*. The dynamics of this phagosomal marker together with the *in silico* data of the MC simulation are in general agreement with previous studies for *D. discoideum* phagosome maturation (Carnell et al., 2011).

Our results on the initial maturation step of acidification in amoebae differ from findings reported for murine and human macrophages infected with *A. fumigatus* conidia. In macrophages, acidification of phagosomes containing was delayed by melanized conidia due to the interference of DHN-melanin with lipid rafts that are required for v-ATPase assembly (Schmidt et al., 2019). The defect in acidification seen for wild-type conidia in macrophages might thus be based on more specific effects on innate immune cells.

The amorphous structure of DHN-melanin and its degradation are still unknown, precluding most biochemical approaches to identify its molecular targets (Nosanchuk et al., 2015). Our *in vitro* results suggest that its degradation might be enhanced by ROS within neutral phagosomes, thereby aggravating its downstream effects on the host cell. Both a neutral to alkaline pH and the presence of ROS have long been known to be key mediators during the biochemical break-down of chemically diverse melanins (Korytowski and Sarna, 1990, Butler and Day, 1998).

Considering the wide environmental occurrence of the fungus, it is probable that DHN-melanin may have additional targets in metazoa when compared to amoebozoan phagocytes. For example, while the MelLec receptor only represents one member of the expanded C-type lectins in metazoa. This family of receptors is restricted to only a few, members in *D. discoideum*. Another possible reason for the difference in acidification of *A. fumigatus* conidia-containing phagosomes in amoebae and macrophages might be due to major differences in the phagosome maturation processes. In *D. discoideum*, we observed that CybA-mCherry, as a proxy for the NOX complex, is delivered to phagosomes at the onset of neutralization, suggesting that v-ATPase has been retrieved at this point. In classically polarized, pro-inflammatory human macrophages (M1), proton pumping and ROS production were found to coincide, thereby maintaining a neutral pH (Canton et al., 2014).

We demonstrated that melanized conidia resided in phagosomes of amoebae for a longer period of time than melanin-deficient conidia and that these phagolysosomes were leaky (Figure 8). Damage to the phagolysosomal membrane might be partially due to the intrinsic production of ROS and might be further enhanced by fungal mycotoxins, such as the spore-borne polyketide trypacidin (Mattern et al., 2015). In *D. discoideum* the ESCRT machinery is effectively recruited to damaged intracellular membranes, such as *Mycobacterium* marinum--containing vacuoles (Cardenal-Munoz et al., 2017, López-Jiménez et al., 2018). In mammalian cells, ESCRT-III recruitment to damaged plasma membranes and lysosomes was hypothesized to depend on the recognition of a local increase of Ca^2+^ by ALIX and/or ALG2 (Jimenez et al., 2014, Skowyra et al., 2018). In *D. discoideum*, ESCRT-III recruitment to sites of membrane damage appears to be independent of Ca^2+^, but strongly depends on Tsg101 (López-Jiménez et al., 2018). While infection with non-melanized conidia ESCRT-III was recruited to phagosomes containing unmelanized conidia, this recruitment did not occur with melanized conidia despite similarly high levels of damage, indicating that this defect in recruitment might be a major cause of proton leakage. How DHN-melanin or its degradation products suppress the cell-autonomous repair machinery of the amoeba host is not clear, but previous observations found that conidia are able to germinate inside certain types of macrophages and amoeba (Schaffner et al., 1982, Van Waeyenberghe et al., 2013, Hillmann et al., 2015). It is likely that damage to the sealed phagolysosome might lead to an influx of nutrients and will thus help the fungus to establish a germination niche. Although this advantage may be restricted to non-specialized phagocytes that are unable to kill the fungus, we also found a protective role for DHN-melanin when encountering a fungivorous amoeba, demonstrating that surface exposure of DHN-melanin provides an overall selective advantage in phagocytic predator-prey interactions in environmental reservoirs.

**Fig 8.**
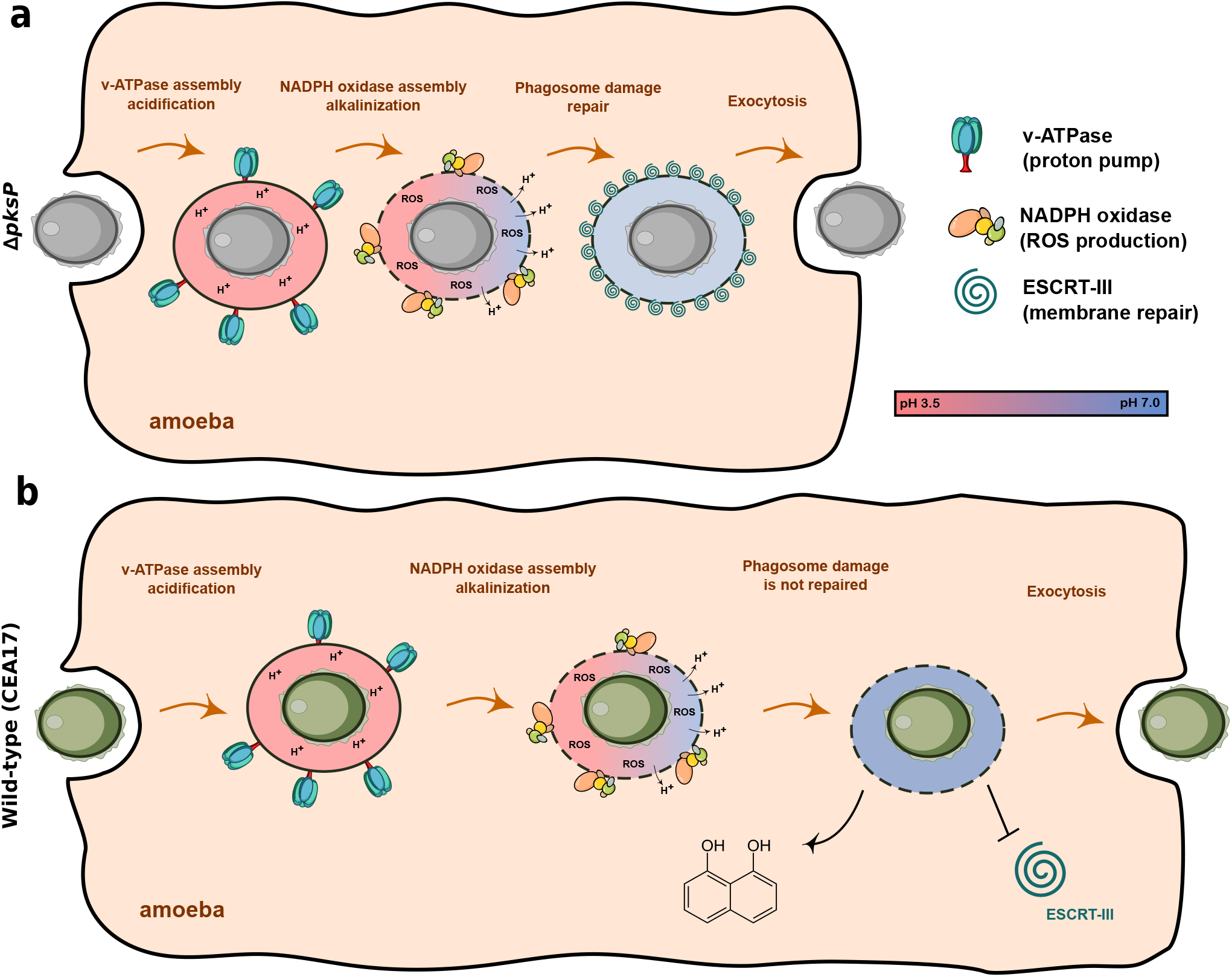
Model for phagosome maturation of *D. discoideum* infected with conidia of *A. fumigatus*. Conidia are acidified in phagosomes within the first minutes after uptake. This process is only marginally affected by DHN-melanin. However, intracellular re-neutralization via ROS is delayed. Intracellular processing induces damage to phagolysosomes which recruits the ESCRT-III repair machinery only with melanin deficient conidia, which are subsequently undergoing exocytosis. The recruitment is repressed by DHN-melanin inducing more damage with prolonged intracellular retention.

## Materials and Methods

### Strains and culture conditions

All strains used in this work are listed in Table S1. *D. discoideum* cells were axenically grown in plastic petri dishes (94×013;16 mm, Greiner Bio-One, Austria) in HL5 medium (Formedium) supplemented with 1 % [w/v] glucose and with 10,000 U g/ml of penicillin and 10 mg/ml of streptomycin (7050218, Genaxxon bioscience) and was split every 2-3 days before reaching confluency. *Protostelium aurantium var. fungivorum* (Hillmann et al., 2018) was grown in PBS (80 g l^−1^ NaCl, 2 g l^−1^ KCl, 26.8 g l^−1^ Na_2_HPO_4_ × 7 H_2_O, 2.4 g l^−1^ KH_2_PO_4_, pH 6.6) with *Rhodotorula mucilaginosa* as a food source at 22°C. *Aspergillus fumigatus* strains were grown in Aspergillus Minimal Medium (AMM) or Czapek-Dox (CZD, Thermo Fisher Scientific, Germany) Medium at 37°C, supplemented with 1.5 % [w/v] agar for growth on solid media.

### Microscopy and image analysis

Microscopy was carried out on an Axio Observer Spinning Disc Confocal Microscope (ZEISS) using ZEN Software 2.6 software. Fluorescent stains and proteins were excited using the 488 nm, 561 nm laser lines. Quantification of fluorescence intensity was performed using ImageJ (https://imagej.nih.gov).

### Resazurin based survival assay after amoeba predation

A total of 1×10^6^ conidia of *A. fumigatus* were placed in 96-well tissue culture in 100μl CZD media. Conidia were confronted with *P. aurantium* directly (resting conidia) or after preincubation at 37°C for 6 h (swollen) MOI 10. *P. aurantium* was collected from pre-cultures on *R. mucilaginosa*. The liquid medium was aspirated from the plate and washed two times with PPB to remove residual yeast cells. Trophozoites were added at prey-predator ratios 10:1, and incubated at 22°C for 18h. Then, the plate was transferred to 37°C for 1h in order to kill the amoeba and induce fungal growth. Resazurin (R7017, Sigma-Aldrich, Taufkirchen, Germany) 0.002% [w/v] was added to quantify the amount of fungal growth to each well in real time as fluorescence, measured in intervals of 30 min over 80 h at λ_ex_ 532 nm/λ_em_ 582 nm using an Infinite M200 Pro fluorescence plate reader (Tecan, Männedorf, Switzerland).

### Measurement of phagosome acidification

*D. discoideum* cells at concentrations of 10^6^ ml^−1^ were axenically grown as an adherent culture in ibidi^®^ dishes (ibidi, Gräfelfing, Germany) in a total volume of 2 ml HL5 supplemented with 1% [w/v] glucose. To synchronize the physiological status of the *D. discoideum* cells, the plates were cooled down to 4°C 10 minutes (before adding conidia) on an ice-cold metal plate. Conidia were stained with FITC and CF594 fluorophores for 10 min and washed two times with PBS. Then, amoebae were infected with conidia at an MOI of 10 and briefly centrifuged (500 rpm, 2min). Excess media was aspirated and a 1% [w/v] agarose sheet was placed on top of cells (1.5×1.5 cm). Then, cells were imaged at 3 to 1 min frame intervals, for up to 4 hrs with a spinning disc confocal system (Axio Observer with a Cell Observer Spinning Disc unit, ZEISS) using the 63 × oil objective. Image processing and quantification of fluorescence intensity was performed with ImageJ. Under infectious conditions, only cells containing conidia were considered for quantification. The GraphPad5 Prism software was used to perform statistical tests and to plot graphs.

### Calibration curve for the acidification measurements

HL5 media supplemented with 1% [w/v] glucose were buffered to pH 3.5 to 8.0. *A. fumigatus* resting conidia of the wild type or Δ*pksP* strain were stored in buffered media. For pH determinations, the integrated density of at least 10 conidia was measured with ImageJ. Then average log of these values were plotted on the calibration curve graph. In order to determine pH on the sample image the integrated density were back calculated from the calibration graph.

### Visualisation of ROS generation in *D. discoideum*

Amoebae were infected in 8-well Ibidi dish with resting conidia of *A. fumigatus* at an MOI of 10. After 2 hours of co-incubation, the superoxide indicator dihydroethidium (DHE, Thermo Fisher Scientific) was added to the wells up to a final concentration of 10 μM. After 10min sample was imaged with a red and blue laser. Experiments were performed in three biological replicates.

### Co-incubation with Dextran

*D. discoideum* cells were incubated with dextran at an MW 70,000 (labelled with RITC, R9379, Sigma-Aldrich), dextran at an MW 4,400 (also labelled with TRITC, T1037, Sigma-Aldrich) and blue dextran MW 5,000 (90008, Sigma-Aldrich). Final concentrations were at 0.5 mg ml^−1^ (70 kDa Dextran), or 1.5 mg ml^−1^, (4.4 and 5 kDa Dextran). After 2 h the cells were washed with fresh media to remove extracellular dextran and were infected with fungal conidia. Dextran loaded lysosomes would fuse with conidia containing phagosomes, thus creating fluorescent rings around the ingested conidia. Damage to the plasma membrane was visible due to selective diffusion of the smaller dextrans into the cytosol.

### Computational modeling

A Monte-Carlo computational model was used to assess the population-wide distribution of acidic phagosomes during infections with conidia of the wild type and Δ*pksP*. Statistical differences were calculated with a Bonferroni post hoc test after two-way ANOVA (P<0.0001). This simulation performs a risk analysis by building models taking into account a range of values obtained in previous experiments (such as phagocytosis rate, the average time of the conidia inside of acidified phagosome, time of exocytosis for each fungal cell line). It then repeatedly executes the calculation, each time using a different set of random values from the probability functions. The generated simulations produce distributions of possible outcomes of the infection for the whole amoeba population for the each fungal cell line. The simulation code is available online at https://github.com/devlxf/FungiSim.

### Synthetic polymerization of 1,8-DHN-melanin

Melanin ghosts were prepared as described by (Youngchim et al., 2004) in concentration of 10^9^ particels ml^−1^. Synthetic melanin was polymerized spontaneously from 1,8-dihydroxynaphthalene (Sigma) over three days in PPB buffer in 48-well plate. Then, H_2_O_2_ were added in various concentrations to the wells. The plates were imaged after two days.

## Acknowledgments

We thank Jean Paul Latgé, Emilia Mellado and Axel A. Brakhage for the supply of *A. fumigatus* strains. This work was supported by a grant of the European Social Fund ESF “Europe for Thuringia” (2015FGR0097 to F.H.) and a grant from the German Research Foundation (DFG, HI 1574/2-1). I.F. was supported by an EMBO Short-Term Fellowship (7020). Work in the T.S. laboratory was supported by a grant from the Swiss National Science Foundation (310030_169386).

## Author Contributions

Conceptualization, I.F., T.S. and F.H.; Methodology, I.F, and J.D.D.; Software, A.F.; Formal Analysis, I.F.; Investigation, I.F.; Resources, T.S. and F.H.; Writing – Original Draft, I.F. and F.H.; Writing – Review & Editing J.D.D., T.S. and F.H.; Visualization, I.F.; Supervision, T.S. and F.H.; Funding Acquisition, I.F., T.S. and F.H.

