## Supplementary Figures and Table for "Conidial melanin of the human pathogenic fungus *Aspergillus fumigatus* disrupts cell autonomous defenses in amoebae"

### Supplementary Figure 1

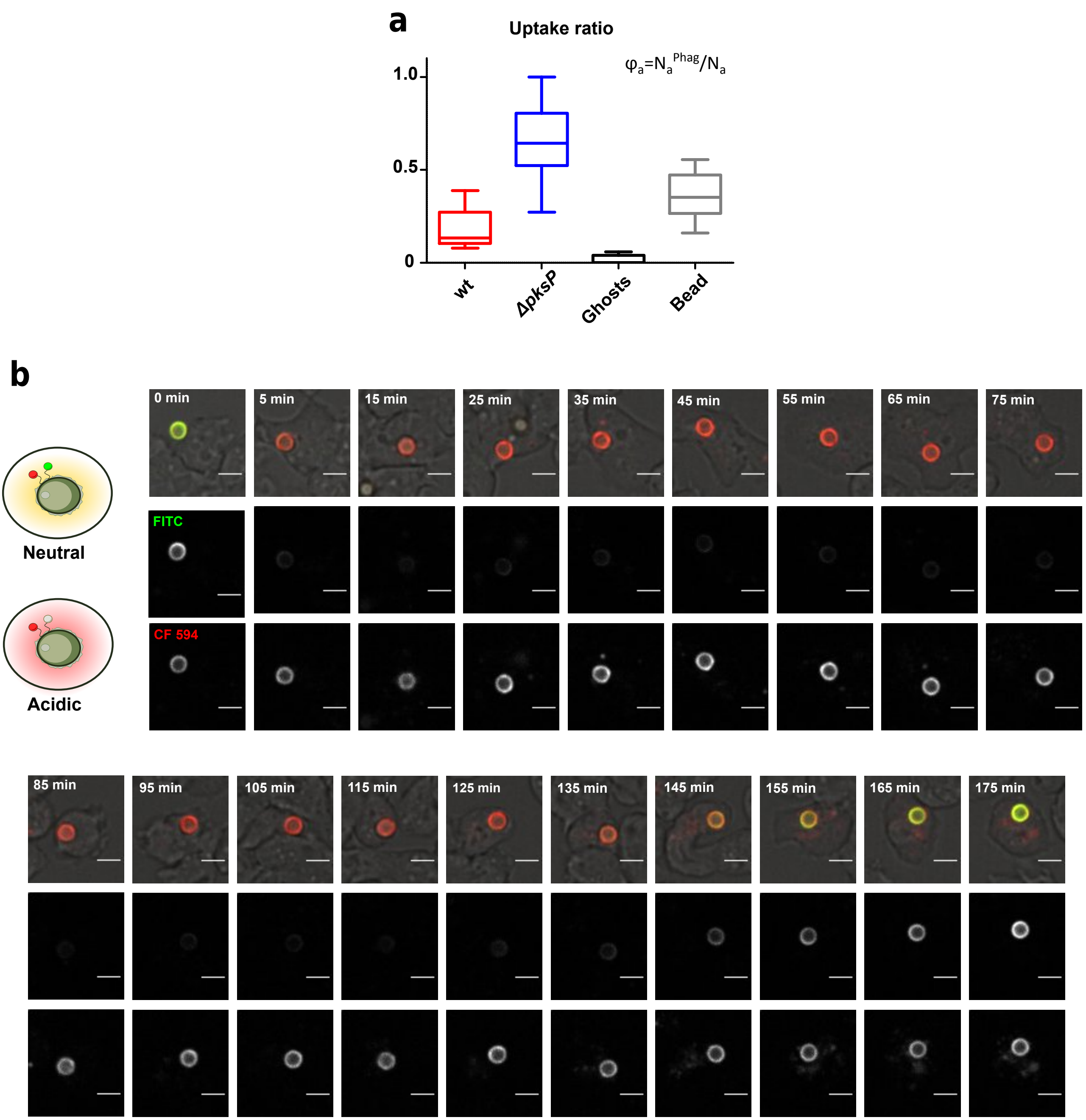


**Supplementary Fig 1.** Phagocytosis efficiency of resting conidia of *A. fumigatus* by *D. discoideum*. a Uptake ratio (amoeba-perspective) **b** A time-lapse illustration of major steps during the phagocytic cycle for resting conidia of the wild type pre-stained with the pH-sensitive fluorophore (FITC) and the reference fluorophore (CF594) to measure acidification and retention time of conidia in *D. discoideum*. The scale bar is 5 μm.

### Supplementary figure 2

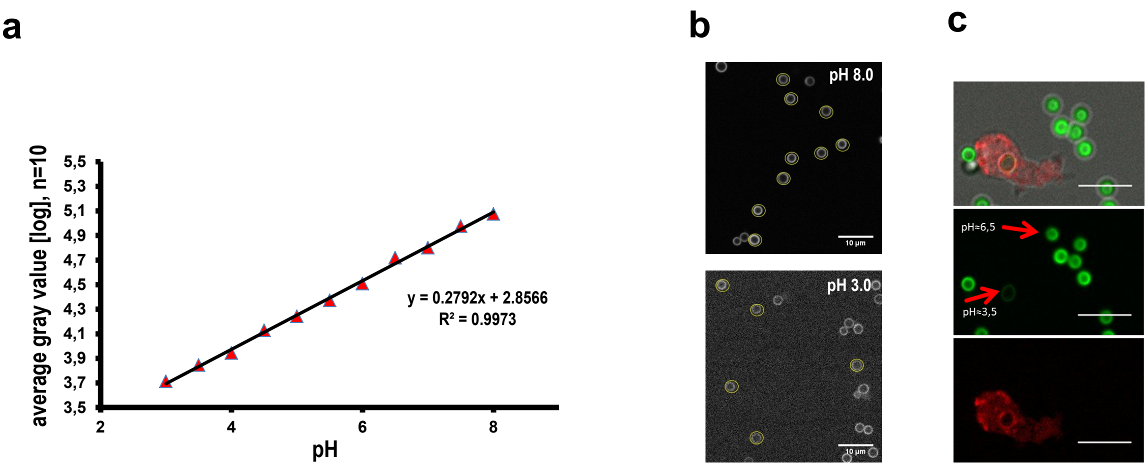


**Supplementary Fig 2.** FITC based pH measurement for conidia of *A. fumigatus*. **a** Example of a calibration curve for the FITC stained fungal conidia imaged at defined pH values. **b** For the determination of the fluorescence intensity 10 random conidia were image in media buffered at different pH. **c** The internalized fungal conidia reside in phagosomes with low pH and assembled VatB-RFP on the surface.

### Supplementary Figure 3

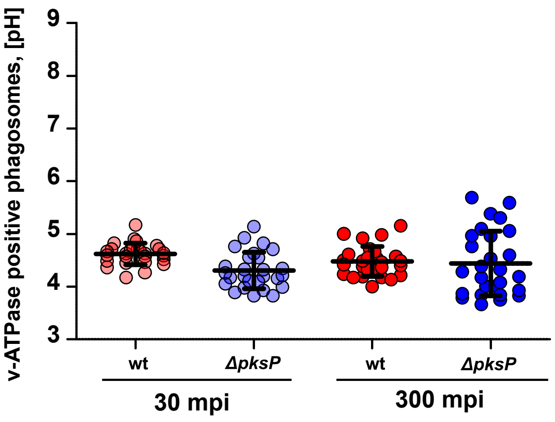


**Supplementary Fig 3.** v-ATPase positive phagosomes are acidic. Measurement of pH inside v-ATPase positive phagosome at different stages of infection. Analyses were performed in three independent experiments and bars indicate the mean and SEM.

### Supplementary Figure 4

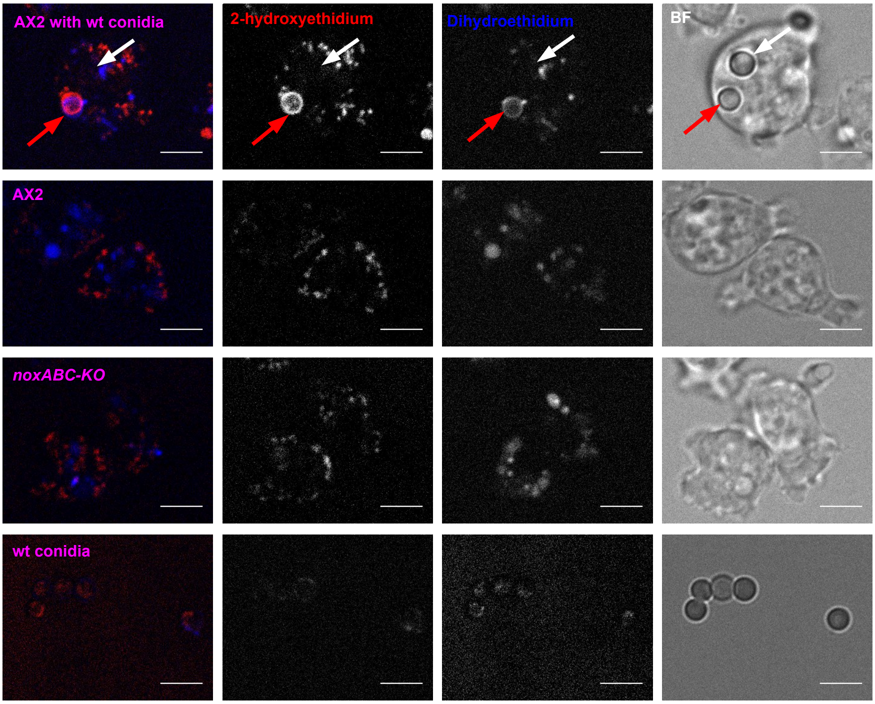


**Supplementary Fig 4.** NADPH oxidase triple knockout *noxABC-KO* of *D. discoideum* mutant generates lower levels of ROS during fungal infection. Amoebae were incubated with fungal conidia for 1h and stained with DHE. Phagosomes containing wild type conidia are either positive (red arrow) or negative (white arrow) for DHE fluorescence.

### Supplementary Figure 5.

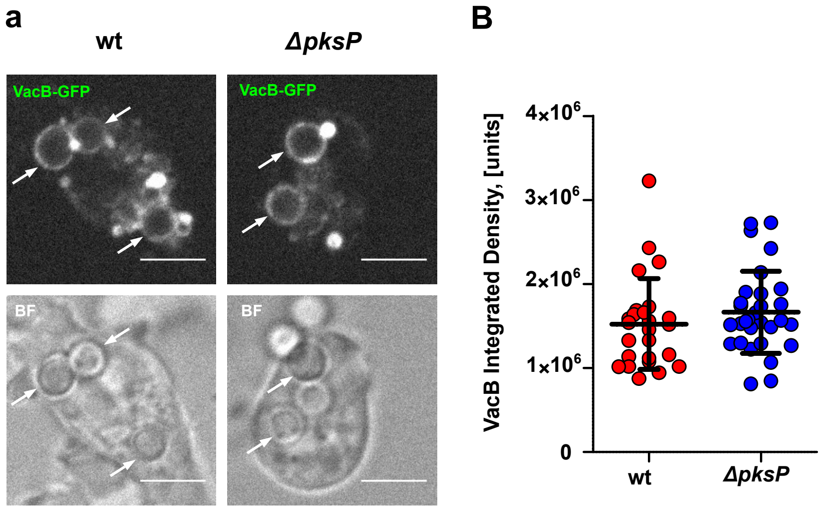


**Supplementary Fig 5.** VacB trafficking on infected phagosomes. **a** Representative images of VacB-GFP-expressing cells infected with conidia of the wild type (wt) or the *ΔpksP* strain at 2 h p.i. White arrows indicate conidia containing phagosomes and colocalization of VacB. Scale bars are 5µm. **b** Quantification of the integrated density for VacB-GFP at conidia containing phagosomes.

### Supplementary Figure 6.

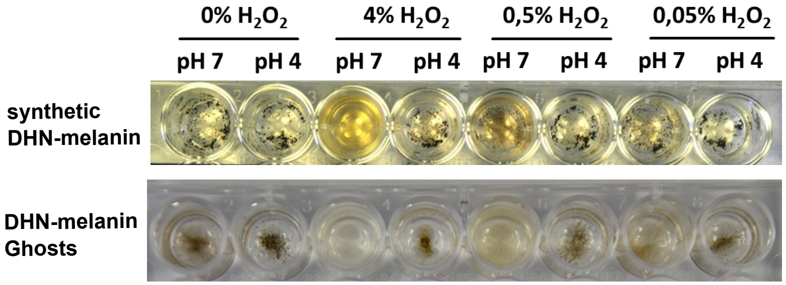


**Supplementary Fig 6.** $H_2O_2$ and pH dependent degradation of synthetic melanin derived from 1,8-DHN or melanin ghosts of wild type conidia of *A. fumigatus*.

### Supplementary Figure 7

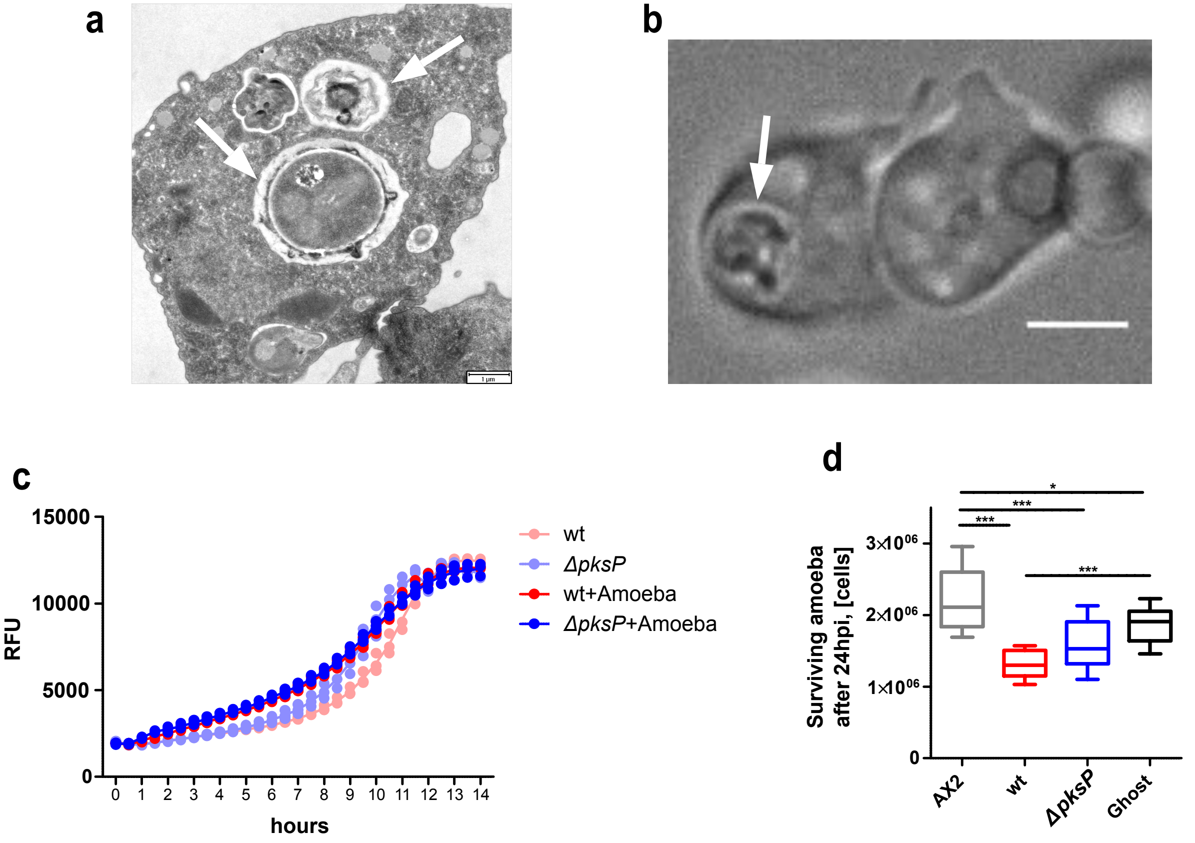


**Supplementary Fig 7.** Viability of swollen conidia of *A. fumigatus* and *D. discoideum* after the confrontation. TEM **a** and light microscopy **b** images of *D. discoideum* containing swollen conidia or conidia-like inclusion at different stages of digestion. **c** Resazurin based measurement of fungal survival after co-incuation with *D. discoideum* at an MOI of 0.01. **d** Viable amoeba cells after 24 h incubation with the fungus. Statistical differences were calculated with a Bonferroni posttest after two-way ANOVA (*p<0.05; **p<0.01; ***p<0.001).

### Supplementary Figure 8

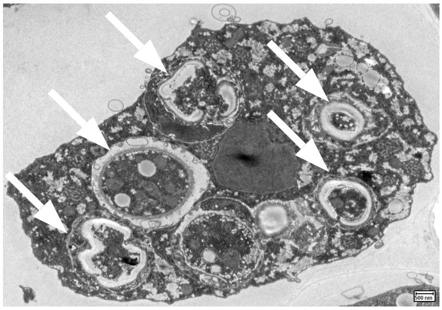

**Supplementary Fig 8.** TEM image of *P. aurantium* with internalized conidia of *A. fumigatus* (white arrows) at different stages of digestion.

**Table S1. Strains used in the study**

***Aspergillus fumigatus***

| Strain | relevant genotype | Background | Resistance | phenotype | Source |
|---|---|---|---|---|---|
| CEA17Δ*akuB* $^{KU80}$ | deletion of Ku family DNA helicase gene | CEA17 | $PyrG^+$ | wild-type | da Silva Ferreira et al., 2006 |
| Δ*pksP* | deletion in the *pksP* gene | CEA17Δ*akuB* $^{KU80}$ | $Hyg^R$ | KO | Hillmann et al., 2015 |
| Δ*rodA* | deletion in the spore coat hydrophobin gene *rodA* | G10 | $Hyg^R$ | KO | Jean Paul Latgé; Thau et al., 1994 |
| G10 | nitrate reductase mutant of strain CBS 144-89 | CBS144.89 | Gln+ | wild-type | Emilia Mellado; Monod et al., 1993 |
| Δ*rodA-47* | deletion in the spore coat hydrophobin gene *rodA* | G10 | $Hyg^R$ | KO | Emilia Mellado |
| Δ*rodA*Δ*rodB-26* | deletion in the spore coat hydrophobin genes *rodA* and *rodB* | Δ*rodA- 47* | $Hyg^R$;$Ble^R$ | KO | Emilia Mellado |
| Δ*rodB-02* | deletion in the spore coat hydrophobin genes *rodB* | G10 | $Ble^R$ | KO | Emilia Mellado |

***Dictyostelium discoideum***

| Strain | | Background | Resistance | | Source |
|---|---|---|---|---|---|
| Ax2 (Ka) | wild-type | | - | wild-type | |
| Ax2 (Gerisch) | wild-type | | - | wild-type | |
| Ax2-NoxABC triple KO | deletion of noxA, B, and C | Ax2 (Gerisch) | Bsr | KO | Zhang et al., 2016 |
| VacB-GFP | *vacB-gfp* (DDB_G0279191) in pDM323 | Ax2 (Ka) | G418 | fluorescent reporter for VacB | Bosmani et al. 2019 |
| VatM-GFP | *vatM-gfp* (DDB_G0291858) in pMJC25 | Ax2 (Ka) | G418 | fluorescent reporter for VatM | Carnell et al., 2011 |
| VatB-RFP | *vatB-rfp* (DDB_G0277401) in pDM451 | Ax2 (Ka) | $Hyg^R$ | fluorescent reporter for VatB | Carnell et al., 2011 |
| GFP-Vps32 | *gfp-vps32* (DDB_G0275573) in pDM317 | Ax2 (Ka) | G418 | fluorescent reporter for Vps32 | López-Jiménez et al., 2018 |
| CybA mCherry | *cybA-mCherry* (DDB_G0267460) | Ax2 (Gerisch) | G418 | fluorescent reporter for CybA | this work |


BOSMANI, C., BACH, F., LEUBA, F., HANNA, N., BURDET, F., PAGNI, M., HAGEDORN, M. & SOLDATI, T. 2019. Dictyostelium discoideum flotillin homologues are essential for phagocytosis and participate in plasma membrane recycling and lysosome biogenesis. bioRxiv, 582049.

CARNELL, M., ZECH, T., CALAMINUS, S. D., URA, S., HAGEDORN, M., JOHNSTON, S. A., MAY, R. C., SOLDATI, T., MACHESKY, L. M. & INSALL, R. H. 2011. Actin polymerization driven by WASH causes V-ATPase retrieval and vesicle neutralization before exocytosis. J Cell Biol, 193, 831-9.

DA SILVA FERREIRA, M. E., KRESS, M. R., SAVOLDI, M., GOLDMAN, M. H., HARTL, A., HEINEKAMP, T., BRAKHAGE, A. A. & GOLDMAN, G. H. 2006. The akuB(KU80) mutant deficient for nonhomologous end joining is a powerful tool for analyzing pathogenicity in Aspergillus fumigatus. Eukaryot Cell, 5, 207-11.

HILLMANN, F., NOVOHRADSKA, S., MATTERN, D. J., FORBERGER, T., HEINEKAMP, T., WESTERMANN, M., WINCKLER, T. & BRAKHAGE, A. A. 2015. Virulence determinants of the human pathogenic fungus Aspergillus fumigatus protect against soil amoeba predation. Environ Microbiol, 17, 2858-69.

LÓPEZ-JIMÉNEZ, A. T., CARDENAL-MUÑOZ, E., LEUBA, F., GERSTENMAIER, L., BARISCH, C., HAGEDORN, M., KING, J. S. & SOLDATI, T. 2019. The ESCRT and autophagy machineries cooperate to repair ESX-1-dependent damage at the Mycobacterium-containing vacuole but have opposite impact on containing the infection. PLOS Pathogens, 14, e1007501.A53+A54

MONOD, M., PARIS, S., SARFATI, J., JATON-OGAY, K., AVE, P. & LATGE, J. P. 1993. Virulence of alkaline protease-deficient mutants of Aspergillus fumigatus. FEMS Microbiol Lett, 106, 39-46.

THAU, N., MONOD, M., CRESTANI, B., ROLLAND, C., TRONCHIN, G., LATGÉ, J. P. & PARIS, S. 1994. rodletless mutants of Aspergillus fumigatus. Infection and Immunity, 62, 4380-4388.

ZHANG, X., ZHUCHENKO, O., KUSPA, A., SOLDATI, T. 2016. Social amoebae trap and kill bacteria by casting DNA nets, Nat Commun. 7, 10938.
